# Lamin-related congenital muscular dystrophy alters mechanical signaling and skeletal muscle growth

**DOI:** 10.1101/2020.08.06.239210

**Authors:** Daniel J. Owens, Julien Messéant, Sophie Moog, Mark Viggars, Arnaud Ferry, Kamel Mamchaoui, Emmanuelle Lacène, Norma Roméro, Astrid Brull, Gisèle Bonne, Gillian Butler-Browne, Catherine Coirault

## Abstract

**Background:** Laminopathies are a clinically heterogeneous group of disorders caused by mutations in the *LMNA* gene, which encodes the nuclear envelope proteins lamins A and C. The most frequent diseases associated with *LMNA* mutations are characterized by skeletal and cardiac involvement, and include autosomal dominant Emery-Dreifuss muscular dystrophy (EDMD), limb-girdle muscular dystrophy type 1B, and *LMNA*-related congenital muscular dystrophy (*LMNA*-CMD). Although the exact pathophysiological mechanisms responsible for *LMNA*-CMD are not yet understood, severe contracture and muscle atrophy suggest that impair skeletal muscle growth may contribute to the disease severity.

**Methods:** We used human muscle stem cells (MuSCs) carrying 4 different *LMNA* mutations and two mouse models of muscle laminopathies, representing a spectrum of disease severity, to investigate the ability of skeletal muscle to differentiate and to hypertrophy in response to mechanical challenges. We extended these finding to individuals with *LMNA*-related muscular dystrophy using muscle biopsies.

**Results:** *In vitro*, we observe impaired myogenic differentiation with disorganized cadherin/β catenin adhesion complexes in MuSCs carrying *LMNA*-CMD. We show that skeletal muscle from *Lmna*-CMD mice is unable to hypertrophy in response to functional overload, due to defective accretion of activated MuSCs, defective protein synthesis and defective remodeling of the neuro-muscular junction. Moreover, stretched myotubes and overloaded muscle fibers with *LMNA*-CMD mutations display aberrant mechanical regulation of the Yes-Associated Protein (YAP), a key sensor and mediator of mechanical cues. We also observe defects in MuSC activation and YAP signaling in muscle biopsies from *LMNA*-CMD patients. These phenotypes are not recapitulated in closely-related EDMD models.

**Conclusions:** Combining studies *in vitro*, *in vivo* and patient samples, we find that *LMNA*-CMD mutations interfere with mechano-signaling pathways in skeletal muscle, implicating defective skeletal muscle growth as a pathogenic contributor for the severity of *LMNA*-related muscular dystrophy.

## INTRODUCTION

Skeletal muscle is a highly organized tissue designed to produce force and movement. It is largely composed of differentiated, multinucleated, post-mitotic myofibers responsible for contraction, and also contains a population of mononucleated muscle stem cells (MuSCs), called satellite cells, that reside between myofibers and the surrounding basal lamina and that display long-term quiescence. Following muscle injury, during post-natal growth and in response to many hypertrophic responses, MuSCs are activated and undergo a highly orchestrated series of events that regulate their proliferation, polarity, and differentiation (reviewed in [1]). Although a subset of MuSCs return to quiescence [2], other activated MuSCs subsequently differentiate and fuse to each other or to existing myofibers. Adhesive contacts between activated MuSCs or between MuSCs and the myofibers are critical to sense and transduce intracellular forces between cells and the extracellular matrix [3, 4] and neighbouring cells [5–7] and provide direct signalling cues essential to stem cell behaviour [8].

Apart from cell adhesive components, recent studies clearly establish that the nucleus is critical for cells to sense and respond to the mechanical properties of their environment [9, 10], thus implicating that muscle plasticity depends on nuclear mechanotransduction. The mechanical properties of the nucleus are largely determined by the nuclear lamina, a fibrous meshwork composed of lamin intermediate filament proteins that underlies the inner nuclear membrane. Nuclear lamins are encoded by three genes: lamin-A and lamin-C (known as A-type lamins) are alternatively spliced products of the *LMNA* gene, whereas lamin-B1 and lamin-B2 (B-type lamins) are encoded by the *LMNB1* and *LMNB2* genes. Mutations in the *LMNA* gene cause laminopathies, a phenotypically diverse group of disorders, including muscular dystrophies and cardiomyopathies [11]. The majority of *LMNA* mutations cause the autosomal dominant Emery-Dreifuss Muscular Dystrophy or EDMD, characterized by progressive muscle wasting, contractures and cardiomyopathy. Lamin-related congenital muscular dystrophy (*LMNA*-CMD) manifests as a particularly severe skeletal muscle phenotype, with muscle wasting beginning very early in life [12], frequent nuclear defects [13] and impaired mechanosensing [14].

Functional loss in A-type lamins alters cytoskeletal actin structures around the nucleus in cells cultured on a rigid substrate [15–17], presumably through an impaired activation of the mechanosensitive transcriptional cofactor serum responsive factor (SRF) and its target genes [18]. *LMNA*-CMD mutations also compromise the ability of cells to adapt their actin cytoskeleton to different cellular microenvironments and to withstand mechanical stretching of the extracellular matrix, owing to the deregulation of Yes-Associated Protein (YAP) signalling pathways [14]. Collectively, these results implicate A-type lamins in modulating the dynamics and organization of the actin cytoskeleton and thus are also implicated in cellular mechanotransduction.

It is currently unknown whether mechanosensing defects in *LMNA*-CMD mutations may explain abnormal skeletal muscle growth seen in laminopathic patients. In the current study, we aim to investigate the role of A-type lamins in the regulation of mechanotransduction at cell-cell adhesions and in multinucleated muscle cells. We also want to determine the consequences of A-type lamin mutations on *in vivo* muscle adaptation to a mechanical challenge. We hypothesize that *LMNA*-CMD mutations impair cellular and molecular mechanisms contributing to skeletal muscle growth. For the first time, we show that *LMNA*-CMD mutations impaired myogenic differentiation *in vitro* due to disorganized cadherin/β-catenin complexes with reduced M-cadherin and β-catenin protein expression. Defective skeletal muscle growth was also revealed *in vivo*, since the *Lmna*-CMD mouse model was unable to hypertrophy due to defective accretion of activated satellite cells and defective protein synthesis in response to functional overload. Moreover, myotubes and muscle fibers with *LMNA*-CMD mutations demonstrate aberrant regulation of YAP nucleo-cytoplasmic translocation in response to different mechanical challenges. More importantly, in a human context, we reported consistent defects in satellite cell activation and YAP signaling in muscle sections from *LMNA*-CMD patients, suggesting that defects in mechano-signaling can contribute to the impaired skeletal muscle growth observed in *LMNA*-CMD patients. Overall, these data strongly suggest that *LMNA*-CMD mutations interfere with satellite cell fate, and as a consequence, skeletal muscle differentiation and growth.

## MATERIELS AND METHODS

### Human myoblasts and cell culture

Muscle biopsies were obtained from the Bank of Tissues for Research (Myobank, a partner in the EU network EuroBioBank) in accordance with European recommendations and French legislation. Written informed consent was obtained from all patients. Experimental protocols were approved by our institution (INSERM). Experiments were performed using immortalized human myoblasts carrying the following heterozygous *LMNA* mutations responsible for *LMNA*-CMD (hereafter referred to as *LMNA*-CMD): a *LMNA* c.94_96delAAG, p.Lys32del (hereafter referred to as ΔK32), *LMNA* p.Arg249Trp (hereafter referred to as R249W), and *LMNA* p.Leu380Ser (hereafter referred to as L380S) mutation [12].

Control immortalized myoblasts were obtained from two healthy control subjects without muscular disorders (hereafter referred to WT1 and WT2). Following muscular biopsy, muscle cell precursors were immortalized and cultured in growth medium consisting of 1 vol 199 Medium /4 vol DMEM (Life technologies, Carlsbad, CA, USA) supplemented with 20% fetal calf serum (Life technologies, Carlsbad, CA, USA), 5 ng/ml hEGF (Life technologies, Carlsbad, CA, USA), 0.5 ng/ml βFGF, 0,1mg/ml Dexamethasone (Sigma-Aldrich, St. Louis, Missouri, USA), 50 µg/ml fetuin (Life technologies, Carlsbad, CA, USA), 5 µg/ml insulin (Life technologies, Carlsbad, CA, USA) and 50 mg/ml Gentamycin (GibcoTM, Life technologies, Carlsbad, CA, USA). Differentiation was induced by switching confluent myoblasts to differentiation medium containing DMEM (Gibco) and 50 mg/ml Gentamycin.

### Immortalized MyoD-converted human myoblasts

EDMD (*LMNA*^H222P^, carrying the heterozygous *LMNA* p.H222P mutation previously described in patient with classical form of EDMD, [19]) and control patient fibroblasts were obtained from skin biopsies and immortalized as previously described [20]. Inducible myogenic conversion was obtained using a doxycycline-inducible Myod1 lentivirus [21]. MyoD-transfected fibroblasts were cultured in a proliferation medium consisting of DMEM, supplemented with 10% fetal bovine serum (Life Technologies) and 0.1% gentamycin (Invitrogen). For myoconversion, doxycycline (2 µg/ml; Sigma Aldrich) was added in the differentiation medium, composed of DMEM with 10 µg/ml Insulin.

### Drug treatments and siRNA

Eukaryotic translation inhibitor Cycloheximide (CHX) (Sigma-Aldrich, St. Louis, Missouri, USA) was diluted in to final concentration of 30 µg/ml in the culture medium and added to adherent myoblasts for 4 hours. siRNA transfections were done with HiPerfect (Qiagen, Venlo, Netherlands) according to manufacturer’s instructions. Downregulation of M-cadherin was observed 72 h after transfection. Sequences of siRNAs are provided in Supplementary Table 1.

### Cyclic strain

Cells were plated on Bioflex culture plates (Flexcell International) coated with fibronectin for 1 day and induced to differentiate for 72 hours. Once myotubes had formed after 72 hours the myotubes were stretched (10% elongation, 0.5 Hz, 4hrs). Following 4hrs stretch, cells were fixed for immunocytochemistry, described below.

### Immunocytochemistry and image analysis

Myotubes were fixed for 5 min with 4% formaldehyde, permeabilized with 0.1%Triton X100 and blocked with 10% bovine serum albumin (BSA) diluted in phosphate buffer solution (PBS). Cells were stained with Phalloidin-Alexa 568 to label F-actin (Interchim, Montluçon, France). The following primary antibodies were used for immunostaining: anti-M-cadherin (Abcam, ab65157), anti-pan-cadherin (Abcam, ab6529), anti-YAP/TAZ (Santa-Cruz, sc-10119s), anti-β catenin (Cell Signaling, cs-9581) and anti-myosin (MF20, DSHB). Secondary antibodies (Life Technologies, Saint-Aubin, France; 1/500) were: Alexa Fluor 488 donkey anti-mouse IgG or Alexa Fluor 488 donkey anti-mouse IgG. Nuclei were stained with Hoechst (ThermoFischer) and Mowiol was used as mounting medium. Confocal images were taken with an Olympus FV 1200 (Olympus, Hamilton, Bermuda) or a laser-scanning microscopy Nikon Ti2 coupled to a Yokogawa CSU-W1 head.

All image analyses were performed using Fiji software (version 1.51). Quantification of β-catenin areas at cell-cell contacts was determined in at least 5 different fields for each experimental condition. For YAP analysis, Z-stacks of images were acquired for each channel, and the middle confocal slice was chosen from the images of the nucleus detected in the Hoechst channel. On the corresponding slice in the YAP channel, the average fluorescence intensity in the nucleus and just outside the nucleus (cytoplasm) was measured to determine the nuclear/cytoplasmic ratio. Fusion index was defined as the number of myosin heavy chain expressing myotubes with greater than 2 nuclei divided by the total number of nuclei.

### SDS-PAGE and protein analysis

Cells were lysed in total protein extraction buffer (50 mM Tris-HCl, pH 7.5, 2% SDS, 250 mM sucrose, 75 mM urea, 1 mM DTT) with added protease inhibitors (25 μg/ml Aprotinin, 10 μg/ml Leupeptin, 1 mM 4-[2-aminoethyl]-benzene sulfonylfluoride hydrochloride and 2mM Na3VO4) or directly in 2x Laemmli buffer. Protein lysates were separated by SDS-PAGE and transferred on PVDF or nitrocellulose membranes. After blocking with bovine serum albumin, membranes were incubated with anti-YAP (Santa-Cruz, sc-10119), anti-M-cadherin (Abcam, ab-65157), anti-β-catenin (cs-9581) or anti-GAPDH (Cell Signaling, cs-2118). Goat anti-mouse, goat anti-rat or donkey anti-goat HRP conjugates were used for HRP-based detection. Detection of adsorbed HRP-coupled secondary antibodies was performed by ECL reaction with Immobilon western chemiluminescent HRP Substrate (Millipore, Billerica, Massachusetts, USA). HRP signals were detected using a CCD-based detection system (Vilber Lourmat) or a G-box system with GeneSnap software (Ozyme, Saint-Quentin, France). Membranes subjected to a second round of immunoblotting were stripped with stripping buffer (62.5mM Tris-HCL pH 6.8, 2%SDS, 100mM β-mercaptoethanol) and incubated at 55°C for 30 minutes with mild shaking before excessive washing with deionized water and re-blocking. Quantification was performed using ImageJ.

### Quantification of gene expression

The mRNA was isolated from cell lysates using the RNeasy mini kit (Qiagen, Hilden, Germany) with the Proteinase K step, according to the manufacturer instruction. The complementary DNA (cDNA) was transcribed by SuperscriptIII (Life Technologies, Carlsbad, CA, USA). Gene expression was quantified by using PerfeCTa-SYBR^®^Green SuperMix (Quanta, Biosciences, Gaithersburg, USA) with the help of LightCycler 480 II (Roche Diagnostics GmbH, Mannheim, Germany). The primers were designed by Primer-BLAST (NCBI) and synthesized by Eurogentec (Liège, Belgium). Expression of all target genes was normalized to the expression of the reference gene *RPLP0*. Primer sequences are listed in Suppl Table 1.

### Animal study

#### Animals

All animal experiments were conducted in accordance with the European Guidelines for the Care and Use of Laboratory Animals and were approved by the institutional ethics committee (APAFIS#2627-2015110616046978). All experiments were performed on male mice. Accredited personnel dedicated to the Care and Use of Experimental Animals has conducted all animal experiments (accreditation numbers #75–1102 and #75–786). *Lmna*^+/ΔK32^ and WT C57Bl/6_129/6J littermates were 3 months of age at the beginning of the experiments. *Lmna*^+/ΔK32^ mice in a C57Bl/6_129/SvJ genetic background were generated by homologous recombination as described previously [22]. The heterozygous *Lmna*^+/ΔK32^ mouse was chosen over homozygous mice as it is the same mutation seen in *LMNA*-CMD patient, increasing the translational potential of the data derived from this model. Additional experiments were performed in the 129S2/SvPasCrl *Lmna*^H222P/H222P^ mice, the mouse model of the classical form of EDMD [23] and compared them with their respective control strains (WT 129S2/SvPasCrl).

#### Functional Overload

*Lmna*^+/ΔK32^, *Lmna*^H222P/H222P^ and their respective control strain mice were used in the study and assigned to overload (FO) or control (CON) groups. Functional overload (FO) of plantaris (PLN) muscles of WT and mutant mice was induced through the tenotomy of soleus and gastrocnemius muscles, in both legs [24]. The muscles were then sutured from the distal tendon to the proximal musculotendinous region leaving the plantaris intact. Animals recovered within 1-2 hours following the end of the procedure and were then monitored daily following surgery for signs of discomfort and infection. For pain management, buprenorphine was administered prior to and following surgery (Vetergesic^©^ 0.3mg/ml, SC: 0.10 mg/kg). At the indicated time (1 and 4 weeks after FO), animals were sacrificed by cervical dislocation and PLN muscles were dissected and processed for molecular or histological analyses. Following removal of visible fat and connective tissue, isolated PLN muscles were quickly frozen in liquid nitrogen cooled isopentane for cryosectioning or snap frozen in liquid nitrogen or fixed in 4% PFA at room temperature for one hour for analysis of the neuromuscular junction and single muscle fibres.

#### In Vivo Estimation of Protein Synthesis

Protein synthesis was measured using the SUnSET method as previously described [25]. In brief, the mice were injected with puromycin (reconstituted in PBS) at a dose of 0.04 µmol.g^−1^ body weight via an intraperitoneal injection exactly 30 min before experimental end point. Muscles were lysed in via mechanical disruption in Roche MagnaLyser tubes containing ceramic beads (Roche, Germany) and ice cold RIPA buffer. Total cell lysate protein content was determined via a BCA protein assay (Pierce, UK). Twenty milligrams of total protein were loaded into a 12% stain-free polyacrylamide gel (BioRad, UK). Electrophoresis was performed for 30 minutes at 250V. Proteins were then transferred onto nitrocellulose membranes. Membranes were blocked for 1 hour with bovine serum albumin (BSA; 5%) then probed for puromycin with a mouse monoclonal puromycin antibody, clone 12D10 (1:20,000 in 5% BSA, Merck Millipore, USA) for 1 hour at room temperature with gentle agitation. The following day, membranes were washed 3×10 minutes with tris-buffered saline-tween (TBS-T). A secondary horseradish peroxidase antibody raised against the same species as the primary antibody was then applied to membranes (1:2000 in 5% BSA, Merck Millipore, USA) for 1 hour at room temperature with gentle agitation. Membranes were washed 3×10 minutes with TBS-T then exposed to a chemiluminescent substrate and imaged on a Bio-Rad Chemi-Doc MP. Total puromycin was calculated relative to total protein.

#### Maximal Force Measures

Maximal isometric tension of the PLN muscle was assessed *in situ* in response to nerve stimulation, as described previously [26]. Briefly, the knee and foot were secured with pins and the distal tendon of the PLN was attached to a lever arm of a servomotor (305B Dual-Mode Lever, Aurora Scientfic) with silk ligature.

#### Immunohistochemistry

Transverse serial sections (8-10 µm) of PLN muscles were obtained using a cryostat, in the mid-belly region. For determination of muscle fiber cross sectional area and minimal Feret diameter, sections were stained with an anti-dystrophin antibody (MANDYS8(8H11) Developmental Studies Hybridoma bank, University of Iowa, USA) to label the myofiber border. Additional sections were stained for laminin (Dako, Z0097), YAP (Santa-Cruz, sc-10119) and/or Pax7 (Developmental Studies Hybridoma bank, University of Iowa, USA). Multiple images were captured of each section using the tile scanning feature on a Leica DM6000 fluorescence wide-field transmission microscope, allowing imaging of the entire section. Myonuclear counts were achieved using an unbiased automated approach; Tile-scanned sections stained for dystrophin and DAPI were coded by one member of the research team and analyzed by Myovision software (www.MyoVision.org) [27] by another member of the team. Myonuclei are defined by the software as any nuclear region having its centroid and greater than 50% of its area inside the sarcolemma.

For the analysis of muscle fiber type, frozen unfixed sections were blocked 1h in PBS plus 2% BSA, 2% sheep serum. Sections were then incubated overnight with primary antibodies against myosin heavy chain (MHC) isoforms (Developmental Studies Hybridoma bank, University of Iowa, USA). After washes in PBS, sections were incubated for 1 h with secondary antibodies (Alexa fluor, Life Technologies, Saint Aubin, France). A minimum of 1500 individual fibers were analyzed per experimental condition. For morphometric analyses, images were captured using a motorized confocal laser-scanning microscope (LSM 700, Carl Zeiss SAS, Le Pecq, France).

Neuromuscular junction (NMJ) analysis was performed on isolated muscle fibres as previously described with minor modifications [28]. Briefly, plantaris muscles were dissected and fixed in 4%PFA/PBS for 1 hour and rinsed with PBS at room temperature. Isolated muscle fibres were washed three times for 15 min in PBS, incubated for 30 min with 100 mM glycine in PBS and rinsed in PBS. Samples were permeabilized and blocked in blocking buffer (4% BSA/5% goat serum/0.5% Triton X-100 in PBS) for 4 hours at room temperature. They were then incubated overnight at 4°C with rabbit polyclonal antibodies against 68 kDa neurofilament (NF, Millipore Bioscience Research Reagents, 1:1000) and synaptophysin (Syn, Thermofisher Scientific, 1:750) in blocking buffer. After four 1-hour washes in PBS, muscles were incubated overnight at 4°C with Cy3-conjugated goat anti-rabbit IgG (Jackson Immunoresearch Laboratories, 1:500) and Alexa Fluor 488-conjugated α-bungarotoxin (α-BTX, Life Technologies, 1:500) in blocking buffer. After four 1-hour washes in PBS, isolated muscle fibres were then flat-mounted in Vectashield (Vector Laboratories) mounting medium. Confocal images were acquired using Leica SPE confocal microscope with a Plan Apo 63x NA 1.4 oil objective (HCX; Leica). Confocal software (LAS AF; Leica) was used for acquisition of Z serial images, with a Plan Apo 63x NA 1.4 oil objective (HCX; Leica). Confocal images presented are single-projected image derived from image stacks. For all imaging, exposure settings were identical between compared samples and groups. Quantifications were done using ImageJ software. AChR cluster area corresponds to the occupied area of α-BTX fluorescent labelling. More than 20 fibres from at least five different mice of each group were analysed.

### Human study

Human muscle sections were obtained from 2 *LMNA*-CMD patients, 1 EDMD patient and 3 control subjects without any muscular disorder. All patients provided informed consent. Clinical summaries and muscle characteristic of all patients are provided in Suppl. Table 2. All the patients underwent an open muscle biopsy for morphological, immunochemical and biochemical analyses on snap-frozen muscle tissue. Transverse serial sections (8-10 µm) were stained with for laminin (Dako, Z0097), YAP (Santa-Cruz, sc-10119) and/or Pax7 (Developmental Studies Hybridoma bank, University of Iowa, USA). Multiple images were captured of each section using the tile scanning feature on a Leica DM6000 fluorescence wide-field transmission microscope, allowing imaging of the entire section.

### Statistical analysis

Graphpad Prism (Graphpad Software, La Jolla, California) was used to calculate and plot mean and standard error of the mean (SEM). Statistical significances were assessed by ANOVA followed by Bonferroni or two-tailed unpaired t-tests. Differences between conditions were considered significant at p < 0.05. Figures were plotted with Graphpad Prism and R with ggplot2 [29].

## RESULTS

### LMNA-CMD muscle stem cells exit the cell cycle but exhibit impaired fusion

We first examined the functional consequences of *LMNA*-CMD mutations on human MuSCs differentiation. To this end, confluent human WT and mutant MuSCs with heterozygous *LMNA* p.Lys32del (*LMNA*^ΔK32^), *LMNA* p.Arg249Trp (*LMNA*^R249W^) and *LMNA* p.Leu380Ser (*LMNA*^L380S^) mutations were shifted from proliferation to differentiation medium (Fig. 1A). We observed a severe reduction in the fusion index in all 3 *LMNA*-CMD mutant cell lines compared with WT (Fig. 1A,B,E). However, *LMNA*-CMD mutated MuSCs were able to arrest cell division and to express myogenin, an early marker for the entry of MuSCs into the differentiation pathway (Fig. 1C,D). No fusion defect was reported in EDMD (*LMNA*^H222P^) mutant cells (Suppl. Fig. 1A,B).

**Figure 1.**
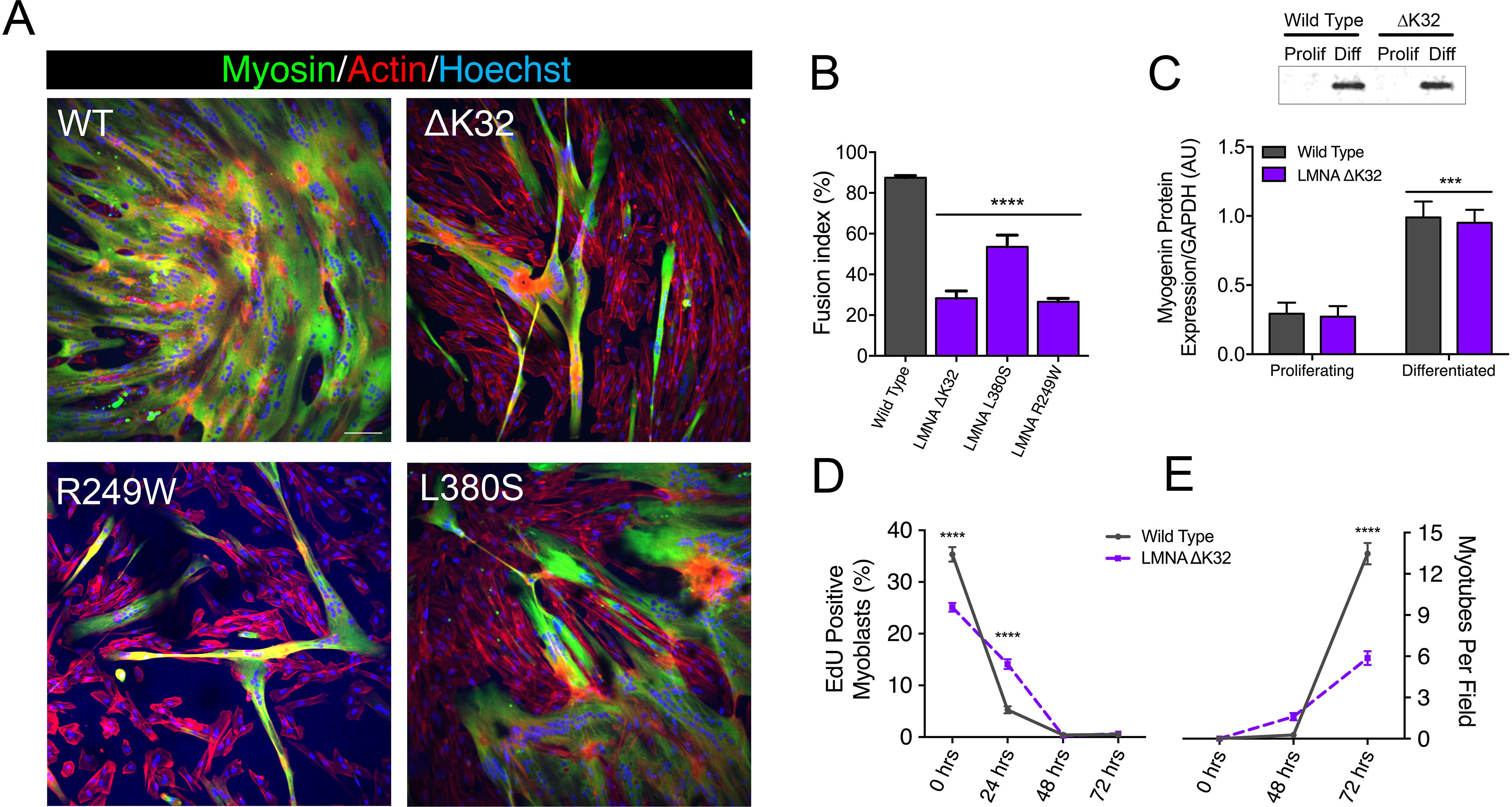
*In vitro* myoblast fusion and myotube formation. **(A)** Confocal immunofluorescence images of myosin (green) in WT and *LMNA*-CMD mutant (ΔK32, L380S and R249W) cells, after 3 days of differentiation. Nuclei are stained with Hoechst (blue). Scale bar= 100 µm. (**B**). Fusion index in WT and *LMNA*-CMD mutant cells after 3 days of differentiation. Pooled values of WT (WT1 and WT2) are presented. Values are expressed as means ± SEM. **** p<0.0001 versus WT myotubes. (**C**) Myogenin expression in WT and *LMNA* ΔK32 mutant cells in proliferation and after 3 days of differentiation. n≥3 from at least 2 separate experiments. *** p<0.001 versus WT myotubes. (**D**) EdU positive myoblasts (%) and (**E**) number of myotubes per field until 3 days of differentiation. Values are expressed as means ± SEM. **** p<0.0001 versus WT cells.

### Impaired cell-cell interactions in LMNA-CMD mutant muscle cells precursors

Fusion defects in *LMNA*-CMD cells prompted us to examine the pattern of cadherin and catenin-based cell adhesion complexes. We immunostained cadherin and β-catenin in confluent MuSCs (Fig. 2A & 3A). In WT MuSCs, cadherin and β-catenin depict the typical “zipper-like” staining pattern at cell-cell contacts, characteristic of force-dependent engagement of the cadherin-catenin complex and cell-cell cohesion (Fig. 2A & 3A). In MuSCs expressing ΔK32, R249W and L380S lamin A/C mutations, both cadherin and β-catenin staining was disorganised with a loss of the “zipper like” staining pattern compared to WT cells (Fig. 2A & 3A). In addition, the size of the β-catenin complex was significantly smaller in *LMNA*-CMD mutant cells compared to WTs (Fig. 3B, each p<0.001). In agreement with the morphological differences, western blot quantification showed that mean protein levels of the muscle-specific cadherin, M-cadherin, and β-catenin were also significantly lower in confluent *LMNA*-CMD mutant MuSCs compared to WT (Fig. 2B & Fig 3C).

**Figure 2.**
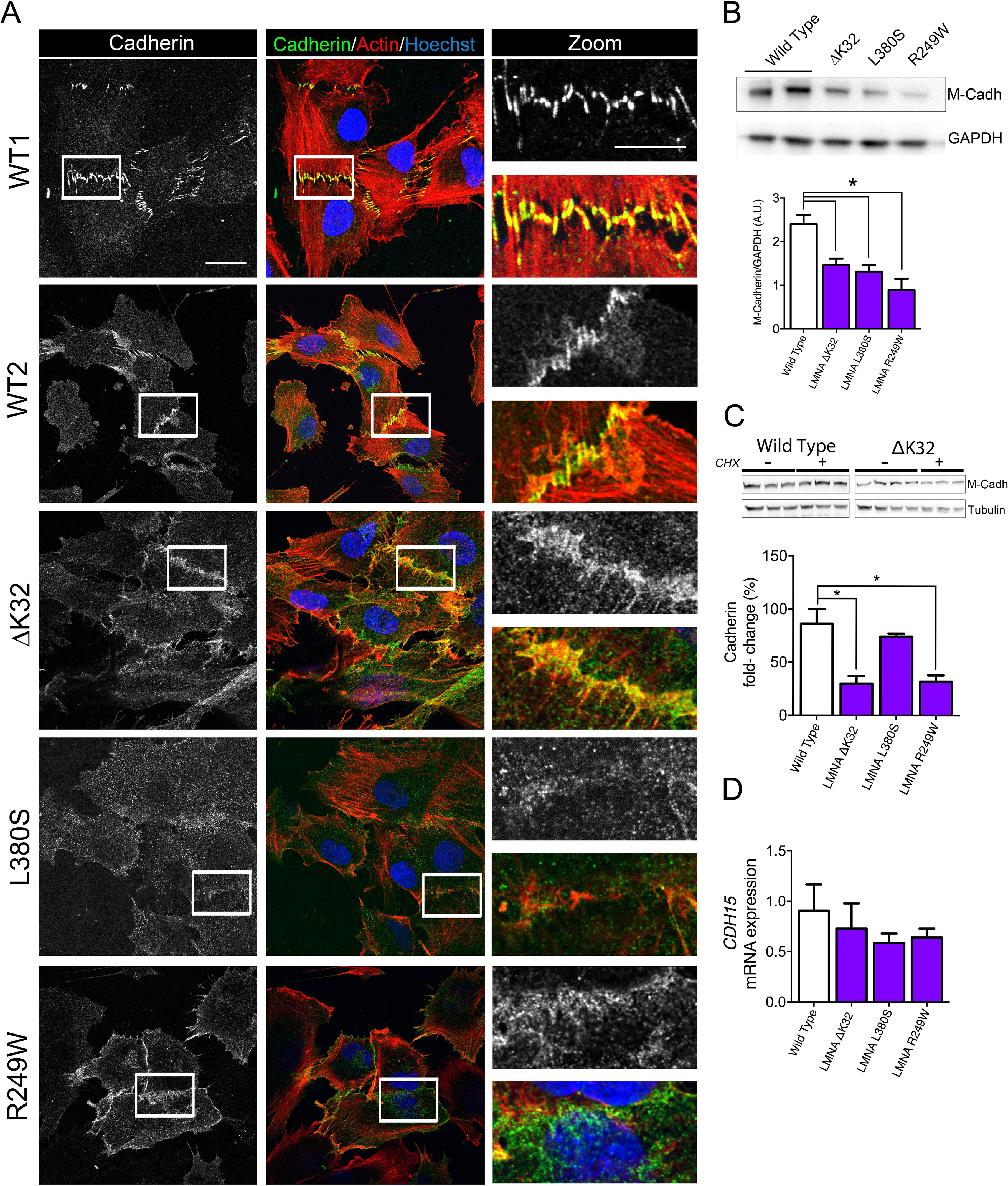
Cadherins in WT and mutant muscle cell precursors. (**A**). Confocal immunofluorescence images of F-actin (phalloidin, red) and cadherin (white or green) in WT (WT1 and WT2) and *LMNA* mutant (ΔK32, L380S and R249W) muscle cell precursors. Nuclei are stained with Hoechst (blue). Scale bar: 20 µm. Zoomed region of cell-cell junctions are shown in left panels. Scale bar: 10µm. (**B**). Top: Representative western blot of M-cadherin and GAPDH expression in WT and *LMNA* mutant myoblasts. Bottom: Quantification of M-cadherin protein levels normalized to GAPDH and expressed in arbitrary units (a.u.). Values are means ± SEM, n≥3 from at least 2 separate experiments. * p<0.005 compared with WT. (**C**) Top: Representative western blot M-cadherin and β-tubulin expression in WT and ΔK32 myoblasts after 4h-treatment with cyclohexamide (CHX). Bottom: Fold-change in M-cadherin protein levels in WT and mutant myoblasts after CHX treatment. M-cadherin protein levels normalized to β-tubulin. Pooled values of WT (WT1 and WT2) are presented. Values are means ± SEM, n=3 in WT and mutant cell lines. * p<0.05 compared with WT. (**D**) mRNA expression of *CDH15* normalized to *RPLP0* and expressed as fold-changes. Pooled values of WT (WT1 and WT2) are presented. Values are means ± SEM, n=3 separate experiments. There was no significant difference between cell lines.

**Figure 3.**
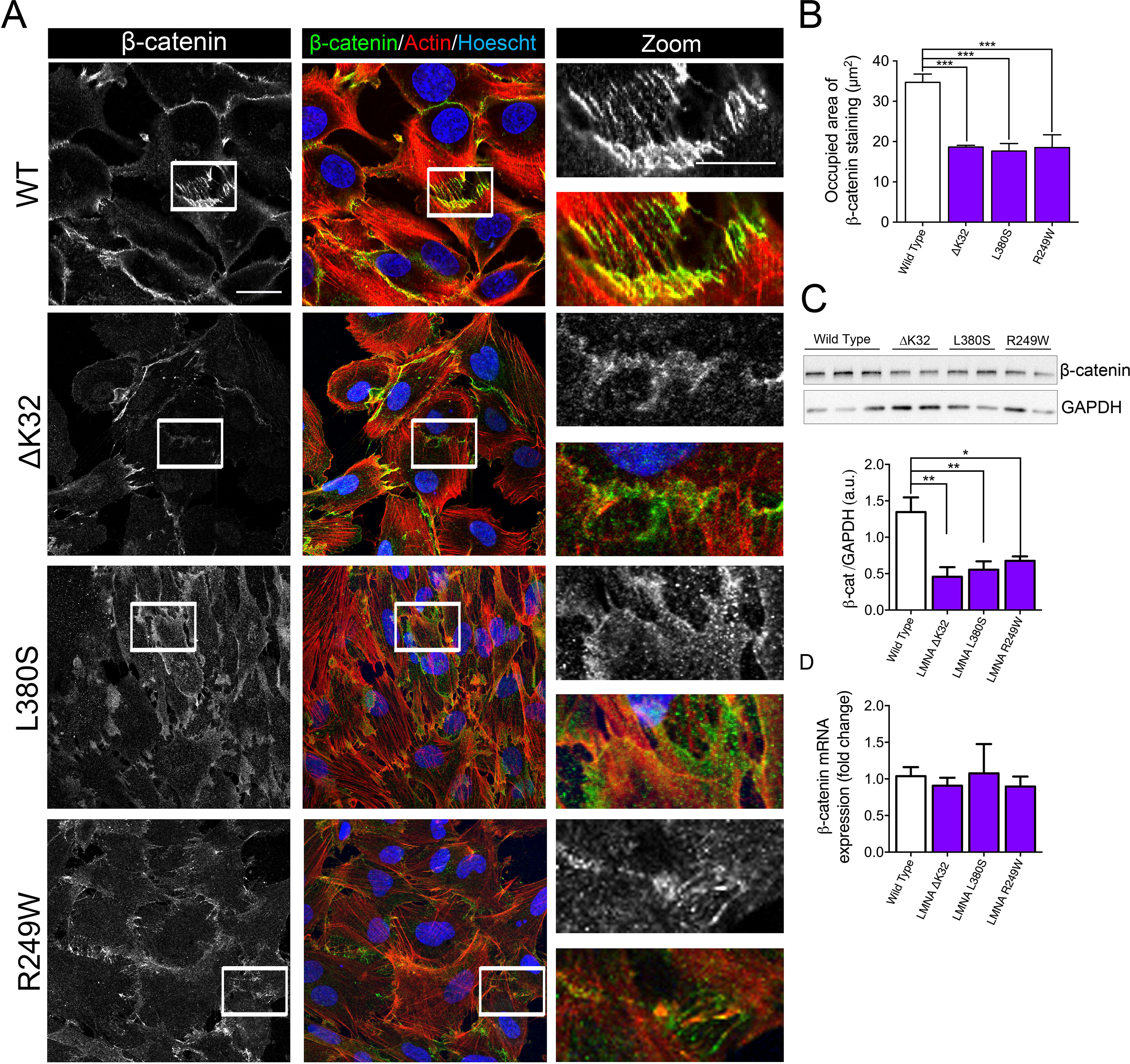
β-catenin in WT and mutant muscle cell precursors. (**A**). Confocal immunofluorescence images of F-actin (phalloidin, red) and β-catenin (white or green) in WT, and *LMNA* mutant (ΔK32, L380S and R249W) mutant myogenic cell precursors. Nuclei are stained with Hoechst (blue). Scale bar: 20 µm. Zoomed region of cell-cell junctions are shown in left panels. Scale bar: 10µm. (**B**). Quantification of the occupied area of β-catenin staining at cell-cell junctions. Pooled values of WT (WT1 and WT2) are presented. Values are means ± SEM from at least 4 different images/cell lines. *** p<0.001 compared with WT. (**C**) Top: Representative western-blot of β-catenin and GAPDH in WT and mutant myoblasts. Bottom: Quantification of β-catenin protein levels expressed in arbitrary units (a.u.). GAPDH was used as a loading control. Pooled values of WT (WT1 and WT2) are presented. Values are means ± SEM, n≥3 from at least 2 separate experiments. * p<0.05, ** p<0.01 compared with WT. (**D**) +

Because the cadherin level is primarily regulated through alteration of its stability [30] we further evaluated M-cadherin degradation in the presence of cycloheximide (CHX), an inhibitor of protein synthesis (Fig. 2C). In confluent *LMNA*-CMD MuSCs, the presence of CHX in the growth medium resulted in a significant reduction of M-cadherin. In contrast, the protein levels of M-cadherin remained relatively constant in WT MuSCs (Fig. 2C). Messenger RNA levels of M-cadherin (*CDH15)* as determined by qRT-PCR were not different between WT and *LMNA*-CMD cells at high cell density (Fig. 2D). These findings suggest that reduced levels of M-cadherin in *LMNA*-CMD mutant MuSCs resulted from an increased degradation of M-cadherin at high cell density and not from reduced transcription. Similarly, β-catenin mRNA expression did not differ between WT and *LMNA*-CMD mutant cells (Fig. 3D). Collectively, these data indicate that the ability to form cadherin/β catenin complexes at cell-cell adhesion sites is impaired in *LMNA*-CMD mutant MuSCs, in part due to degradation of M-cadherin.

### YAP nuclear sequestration in LMNA-CMD mutant myotubes

In non-muscle cells, cadherin bound β-catenin at cell-cell contacts is a critical regulator of YAP localization [31, 32]. YAP nuclear localization promotes the proliferation of MuSCs whilst inhibiting myogenic differentiation [33]. In static WT myotubes, YAP was predominantly cytoplasmic, as previously reported [33]. In contrast, myotubes with *LMNA*-CMD mutations had a predominant nuclear YAP localization, attesting to a defective localization of YAP (Fig. 4A–B). To test whether YAP mislocalization in *LMNA*-CMD was due to reduced M-cadherin protein expression, we treated confluent WT cells with small interfering RNA (siRNA) against M-cadherin and analysed YAP localization (Fig. 4, C, D). Depletion of M-cadherin impaired cell-density dependent redistribution of YAP to the cytoplasm, as attested by significantly higher YAP nucleo-cytoplasmic ratio in cadherin-depleted WT cells (Fig. 4D). These results indicate that reduced M-cadherin-/β catenin mediated cell adherence may impair YAP nuclear localization, and thus interfere with mechanical feedback signalling. Overall, our data strongly suggested that impaired myogenic differentiation in *LMNA*-CMD mutations could be due to impaired interactions between MuSCs.

**Figure 4.**
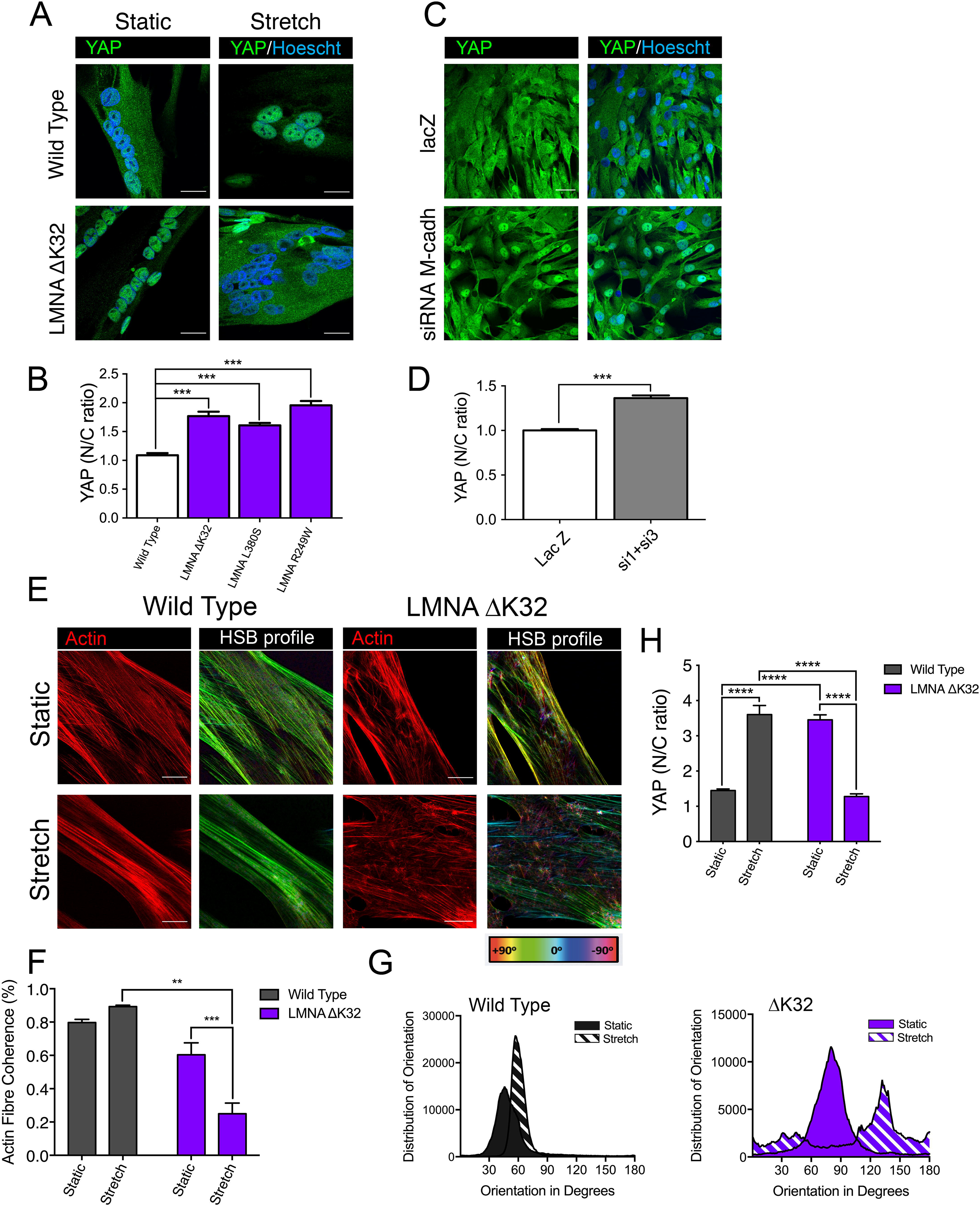
Adaptability of WT and *LMNA* ΔK32 myotubes to cyclic stretch. **(A)** Confocal immunofluorescence images of YAP (green) in WT and *LMNA* ΔK32 mutant myotubes (72h differentiation) in static and after stretch. Nuclei are stained with Hoechst (blue). Scale bar: 20 µm (**B**) Quantification of YAP nucleo-cytoplasmic (N/C) ratio determined by immunocytochemical image analysis in WT and *LMNA* mutant myotubes. Pooled values of WT (WT1 and WT2) are presented. Values are expressed as means ± SEM, n≥60 cell in each cell line. *** p<0.001 compared with WT. (**C**) Immunofluorescence images of YAP (green) in WT treated with lacZ or siRNA against cadherin. Nuclei are stained with Hoechst (blue). Scale bar: 30 µm. **(D)** Quantification of YAP nucleo-cytoplasmic (N/C) ratio determined by immunocytochemical image analysis in WT treated with lacZ or siRNA against cadherin. Values are expressed as means ± SEM, n≥180 cells in each group. *p<0.001 compared with lacZ-treated cells. (**E**) Confocal immunofluorescence images of actin (red) and HSB profile in WT and *LMNA* ΔK32 mutant myotubes in static and after stretch. Nuclei are stained with Hoechst (blue). Scale bar: 20 µm. (**F**) Actin fiber coherence in the dominant direction as determined by analysis of confocal images of myotubes stained fluorescently for actin (phalloidin) in ImageJ using OrientationJ plug-in. N=3 per condition. ** p<0.01, *** p<0.001, compared with WT. (**G**) Representative distribution of actin filaments as a function of the orientation in degrees in static and after stretch. (**H**) Quantification of YAP nucleo-cytoplasmic (N/C) after cyclic stretch in myotubes. Pooled values of WT (WT1 and WT2) are presented. Values are expressed as means ± SEM, n≥65 cell in each cell line. **** p<0.0001 versus control values.

### Adaptability to mechanical constraints is severely affected in LMNA-CMD myotubes

We also found that whilst WT myotubes were able to respond to mechanical stretch by reorganizing the actin cytoskeleton into parallel actin fibers, the cyclically stretched *LMNA*-CMD mutant myotubes displayed actin fibres that lacked orientation (Fig. 4E–G). In addition, cyclic stretching induced relocalization of YAP into the nucleus in WT myotubes whilst nuclear YAP exclusion was observed in stretched *LMNA*-CMD mutant myotubes (Fig. 4A–H). These data suggest that myotubes carrying *LMNA*-CMD mutations are unable to adapt to acute mechanical stretch and appropriately regulate the putative mechanical signalling molecule, YAP that regulates and is regulated by the actin cytoskeleton.

### Defective muscle hypertrophy in Lmna ΔK32 heterozygous mice

We next examined the *in vivo* physiological implications of *LMNA*-CMD-induced muscle defects to sense and respond to mechanical stress. Following functional overload (FO) we found that the plantaris muscle (PLN) from *Lmna*^+/ΔK32^ mice hypertrophied significantly less than their WT counterparts (Fig. 5A). Muscle mass normalized to body mass was comparable at baseline (WT = 0.69±0.02 mg.g^−1^ vs *Lmna*^+/ΔK32^ = 0.65±0.02 mg.g^−1^). However, normalized PLN mass significantly increased in WT mice after 7 days (1.15±0.07 mg.g^−1^) and 4 weeks (1.59±0.06 mg.g^−1^) of overload, whereas mutant mice showed a significantly impaired adaptive response at both 7 days (0.84±0.05 mg.g^−1^) and 4 weeks (1.27±0.06 mg.g^−1^). In addition, FO significantly increased nuclear deformability in *Lmna*^+/ΔK32^ (Suppl. Fig. 2A,B), thus validating *in vivo* the increased nuclear deformability previously reported *in vitro* [13]. WT mice also showed significantly improved maximal force production of PLN in response to 4 weeks of overload compared to mutant mice (p<0.05; Fig. 5B). In contrast, muscle of *Lmna*^H222P^ mice were able to respond to overload to a similar extent to their WT counterparts (Suppl. Fig. 1C), suggesting that impaired mechano-sensitivity and muscle growth are specific to congenital laminopathies.

**Figure 5.**
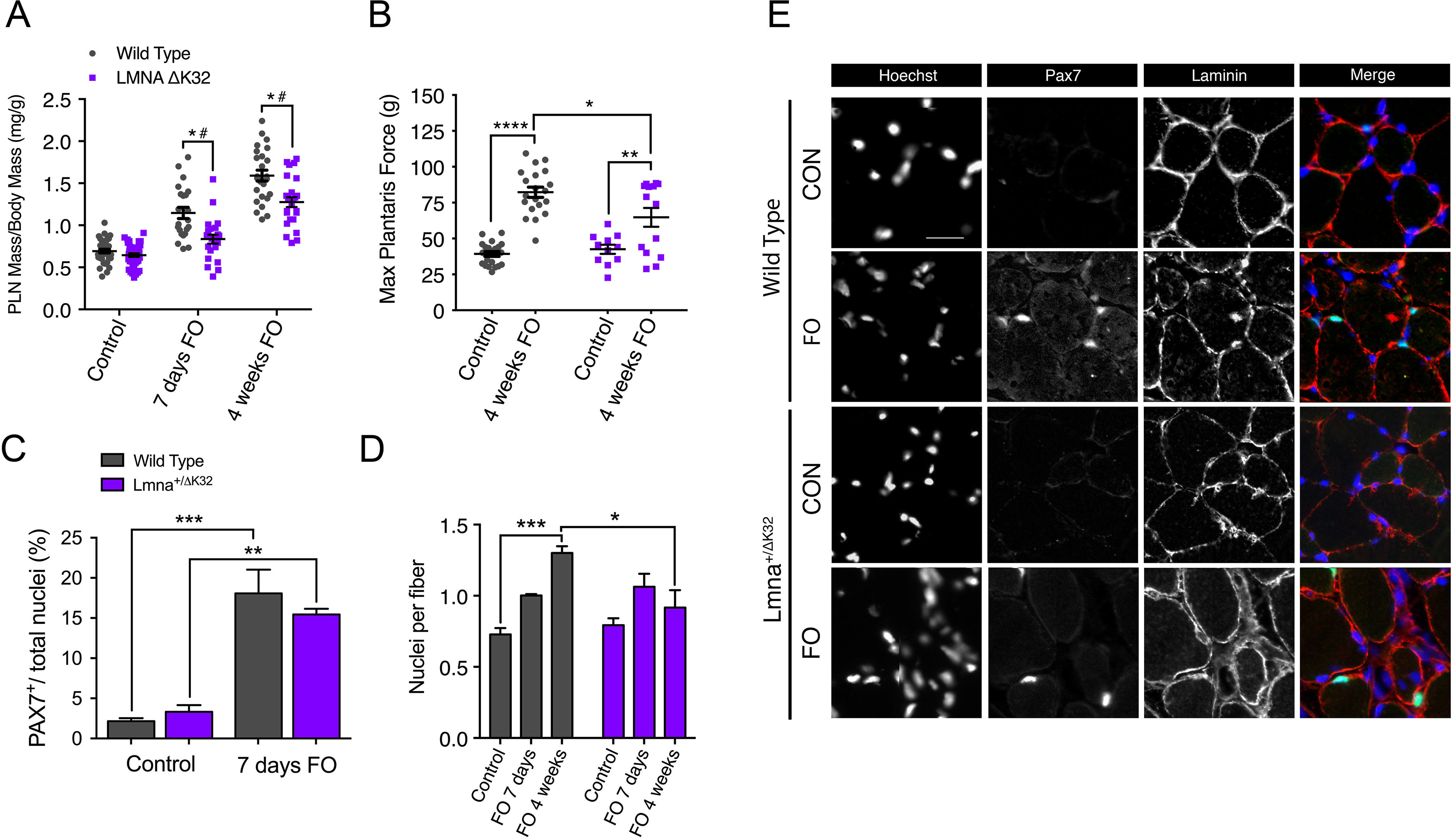
Functional and morphological abnormalities of *Lmna*^+/ΔK32^ mice to functional overload. (**A**) Plantaris muscle mass normalized by body mass from WT and *Lmna*^+/ΔK32^ mice in control and after 7-days and 4-weeks FO. **** p<0.0001 versus WT and versus control conditions. (**B**) Plantaris muscle maximal force from WT and *Lmna*^+/ΔK32^ mice in control and after 4-weeks FO. * p<0.05 and **** p<0.001 versus control conditions. (**C**) Quantification of Pax7^+^ cells as a percentage of total nuclei in control and after 7-days FO. Values are expressed as means ± SEM, *p<0.05 versus control condition. (**D**) Quantification of nuclei per fiber from WT and *Lmna*^+/ΔK32^ mice in control and after 7-days and 4-weeks FO as determined by quantification of Hoechst stained whole tissue sections by Myovision software. ** p<0.005 and *** p<0.001. (E) Immunofluorescence images of PAX7^+^ (green) and laminin (red) in plantaris muscle section in WT and Lmna+/ΔK32 mice in control and after 7-days FO. Nuclei are stained with Hoechst (blue). Scale bar: 25 µm.

### Defective myonuclear accretion in Lmna ΔK32 heterozygous mice

Myonuclear accretion is thought to be a determinant of exercise-induced remodelling in skeletal muscle [34] and myonuclear accretion relies on the activation and proliferation of MuSCs, and fusion of the activated MuSCs into new and existing myofibers [35]. MuSC fusion involves the formation of cell-cell contacts, a process regulated by cell-cell adhesion molecules β-catenin and M-cadherin, which we found to be dysregulated in *LMNA*-CMD mutant cells *in vitro* (Figures 2,3). We therefore examined whether MuSC’s from *Lmna*^+/ΔK32^ mice could be activated and incorporated into new and existing myofibers in response to functional overload (FO). After 1 week of FO, both WT and *Lmna*^+/ΔK32^ mutant mice had an increased number of Pax 7+ cells (Fig. 5C,E) indicating MuSCs proliferation. To determine the fusion capacities of Pax 7^+^ cells, the number of myonuclei (Hoechst staining) inside the sarcolemma (dystrophin immunostaining) were counted in control and mutant PLN muscle sections before and at 1 and 4 weeks after FO. The number of myonuclei per myofiber was similar before FO between WT and mutant muscles and increased significantly in WT PLN muscles following 4-weeks of FO (Fig. 5D). Conversely, myonuclei number did not change in *Lmna*^+/ΔK32^ mice at 1 and 4 weeks of FO and was significantly lower than in WT after 4-weeks of FO (Fig. 5D). Taken together, our data show that *Lmna*^+/ΔK32^ mutation leads to a lack of myonuclear accretion in response to functional overload, despite activation and proliferation of MuSCs.

### YAP abundance is higher at baseline but decreases after FO in Lmna ΔK32 heterozygous mice

To assess whether defective hypertrophy could be a consequence of aberrant mechano-responsiveness in mature fibers and muscle precursor cells, we analysed YAP signalling in PLN from *Lmna*^+/ΔK32^ mice. It is well established that YAP stimulates muscle fiber hypertrophy and protein synthesis [36, 37]. In control conditions, YAP labelling was clearly detectable at the muscle fiber membrane and was also detected in some nuclei localized within the laminin boundary of muscle fibers (Fig. 6A), consistent with previous data [36, 37]. However, before FO, the number of YAP^+^ fibers was significantly higher in *Lmna*^+/ΔK32^ compared to WT mice (p<0.001, Fig. 6A,B. After 1-week FO, the number of YAP^+^ fibres significantly increased in the WT mice whilst YAP staining was markedly reduced in *Lmna*^+/ΔK32^ mice (Fig. 6A,B). Four weeks after the FO procedure, the number of YAP^+^ fibers had returned to baseline in the WT (Fig. 6A,B). In the *Lmna*^+/ΔK32^ mice, YAP was still downregulated after 4-wk FO compared to corresponding baseline values.

**Figure 6.**
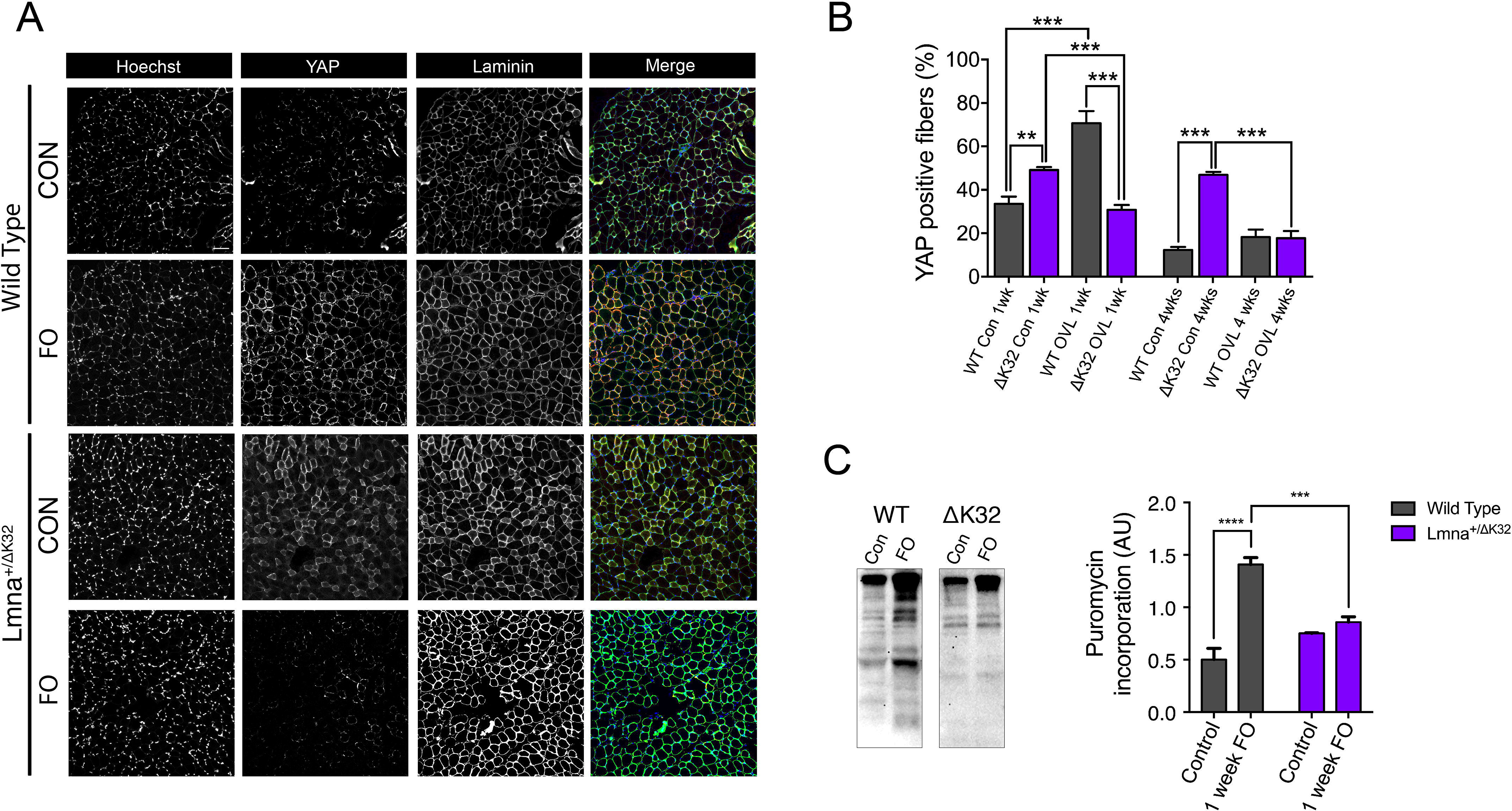
(**A**) Immunofluorescence images of YAP (green) and laminin (red) in control and 7-days FO plantaris muscles from WT and *Lmna*^+/ΔK32^ mice. Nuclei are stained with Hoechst (blue). Scale bar: 100 µm. (**B**) Quantification of YAP^+^ fibers in control and 7-days FO plantaris muscles from WT and *Lmna*^+/ΔK32^ mice. ** p<0.01 versus WT, *** p<0.001 versus control conditions. (**C**) Representative western blot and quantification of puromycin incorporation in control and 7-days FO plantaris muscles from WT and *Lmna*^+/ΔK32^ mice. *** p<0.001 versus WT. Values are expressed as means ± SEM

### Defective muscle protein synthesis in Lmna ΔK32 heterozygous mice

We reasoned that the increase in YAP signaling seen in the early stages of remodelling in overloaded WT muscle, would be associated with increased rates of protein synthesis. By employing the SUnSET method which measures acute incorporation of puromycin into newly synthesised peptides [25], we determined the rate of protein synthesis in PLN muscle of CON and FO mice after 7 days of FO. As hypothesised, PL of WT mice responded to FO by increasing rates of protein synthesis (P<0.0001), whereas *Lmna*^+/ΔK32^ mice did not (P=0.0344; Fig. 6C).

### Neuromuscular junction defects after functional overload in Lmna ΔK32 heterozygous mice

Since the neuromuscular junction (NMJ) is an adaptable/plastic synapse, highly sensitive to decreased or increased activity, we next decided to analyse morphological changes occurring at NMJs of WT and *Lmna*^+/ΔK32^ mice following 4 weeks of FO. Muscle fibers were isolated from plantaris muscle and stained with α-bungarotoxin (α-BTX) to label acetylcholine receptor (AChR) clusters as well as antibodies against neurofilament (NF) and synaptophysin (Syn) to visualize axon and nerve terminals respectively (Fig. 7). In control condition, both WT and *Lmna*^+/ΔK32^ mice exhibited mature NMJs characterized by an elaborate continuous topology that have a pretzel-like shape (Fig. 7A). Interestingly, we observed a significant increase in AChR clusters area both in WT (p<0.0005) and *Lmna*^+/ΔK32^ (p<0.05) mice after FO, as determined by fluorescent labelling with α-BTX (Fig.7Bi). The nerve terminal area of both WT and *Lmna*^+/ΔK32^ was also significantly increased following FO and pre-post synaptic overlap was unaffected by FO in both strains of mice (Fig. 7B[ii] and [iii]). However, despite *Lmna*^+/ΔK32^ demonstrating an increase in AChR cluster area in response to FO, postsynaptic architecture appeared discontinuous with isolated AChR clusters (Fig. 7A). Morphometric analysis revealed that the number of AChR clusters per NMJ (ie. the number of continuous AChR-stained structures per synapse) was significantly increased in *Lmna*^+/ΔK32^ following FO indicating a severe dismantlement of the postsynaptic counterpart (Fig. 7B[iv]). These results suggest NMJ stability is impaired in *Lmna*^+/ΔK32^ mice following FO.

**Figure 7.**
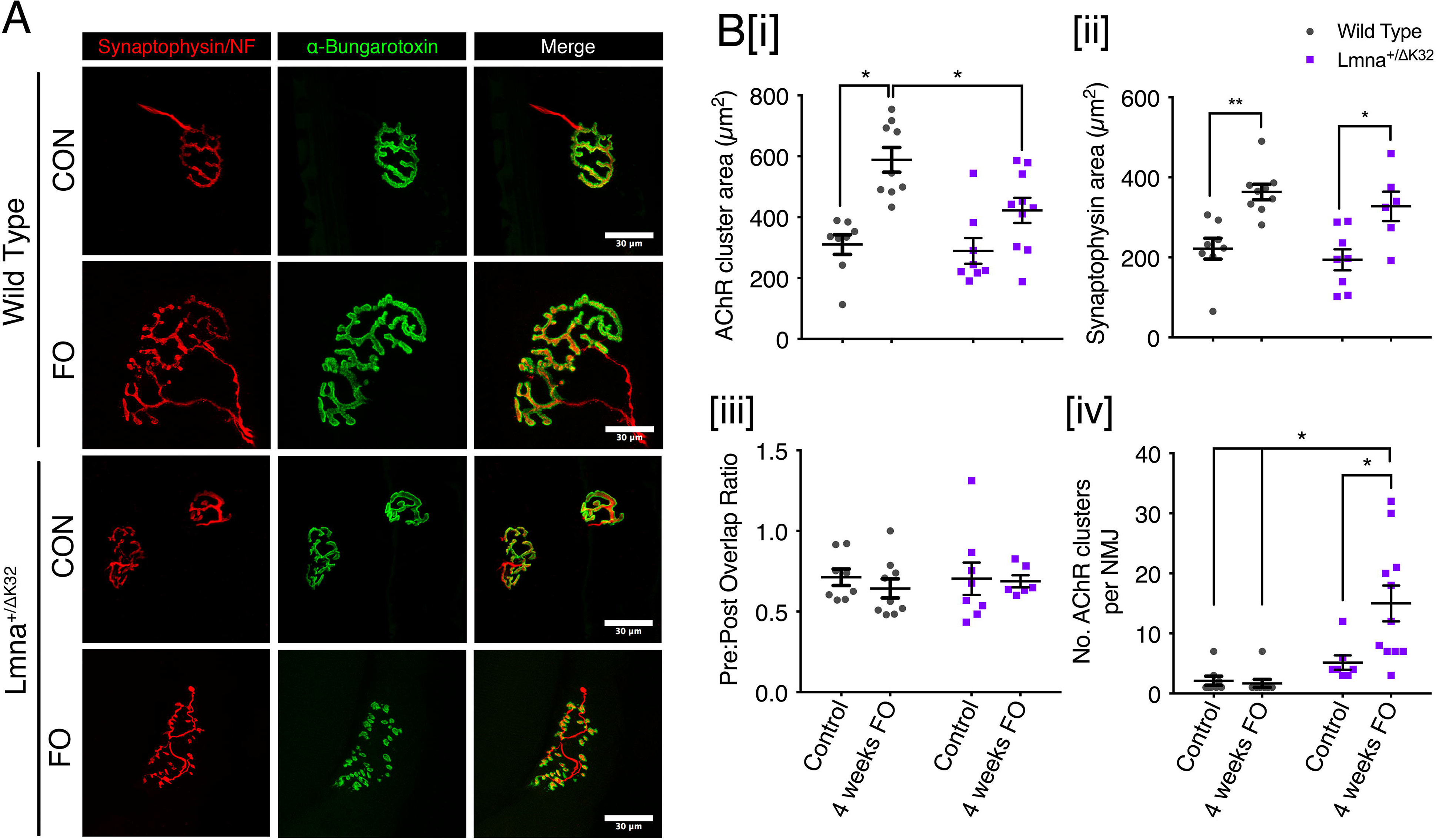
Neuromuscular junction defects of *Lmna*^+/ΔK32^ mice following functional overload. (**A**) Confocal immunofluorescence images of pre-synaptic structure (synaptophysin/neurofilament; red), post synaptic structure (α-bungarotoxin; green) and merged image. Scale bar: 30 µm. Values are expressed as means ± SEM (**B)[i]** Acetylcholine receptor cluster area in WT and *Lmna*^+/ΔK32^ mice in control conditions and following FO. * p<0.05. (**ii**) synaptophysin area in WT and *Lmna*^+/ΔK32^ mice in control conditions and following FO. * p<0.05. (**iv**) Pre/post synapse overlap (i.e. synaptophysin/α-bungarotoxin) and (**iii**) number of acetylcholine receptor clusters per neuromuscular junction * p<0.05.

### Muscle biopsies from LMNA-CMD patients revealed increased Pax7^+^ cells and YAP signalling

Finally, to corroborate our findings from the *in vitro* patient cells and *in vivo* mouse models of striated muscle *Lmna*-CMD in a clinically relevant context, we examined skeletal muscle biopsy samples from patients and control donors. Satellite cells expressing the Pax7^+^ cells are known to decline with age, we examined the number of Pax7^+^ cells in an age-dependent context. The number of Pax7^+^ cells was 3-to 4-fold higher in the 2 children with *LMNA*-CMD compared to the age-matched control (p<0.001), and nearly 1.5-fold higher to that observed in a control new-born (4 day-old) (Fig. 8A–C and Suppl Fig. 3C). More importantly, almost half of Pax7^+^ cells from human *LMNA*-CMD did not reside in the satellite cell position but instead were found in the interstitial space (Fig. 8C). These data suggest an increased proliferation of MuSCs in *LMNA*-CMD patient biopsies with a decreased ability to fuse. In addition, muscle cryosections were immunofluorescently labelled for YAP. In the control muscle, YAP labelling was clearly detectable in 42±3 % fibers and 13±1 % nuclei localized within the laminin boundary of muscle fibers (Fig. 8D–F). Interestingly, the percentage of YAP^+^ fibers and YAP^+^ nuclei were respectively 2- and 3-fold higher in the *LMNA*-CMD patients compared with their relative age-matched control (each p<0.01), and slightly lower to what observed in a control new-born (Suppl. Fig. 3). Taken together, these data highlight a novel mechanism by which defective accretion of activated MuSCs and impaired YAP signalling contribute to the defect in muscle growth and severe muscle weakness observed in the most severe forms of the human striated muscle laminopathies.

**Figure 8.**
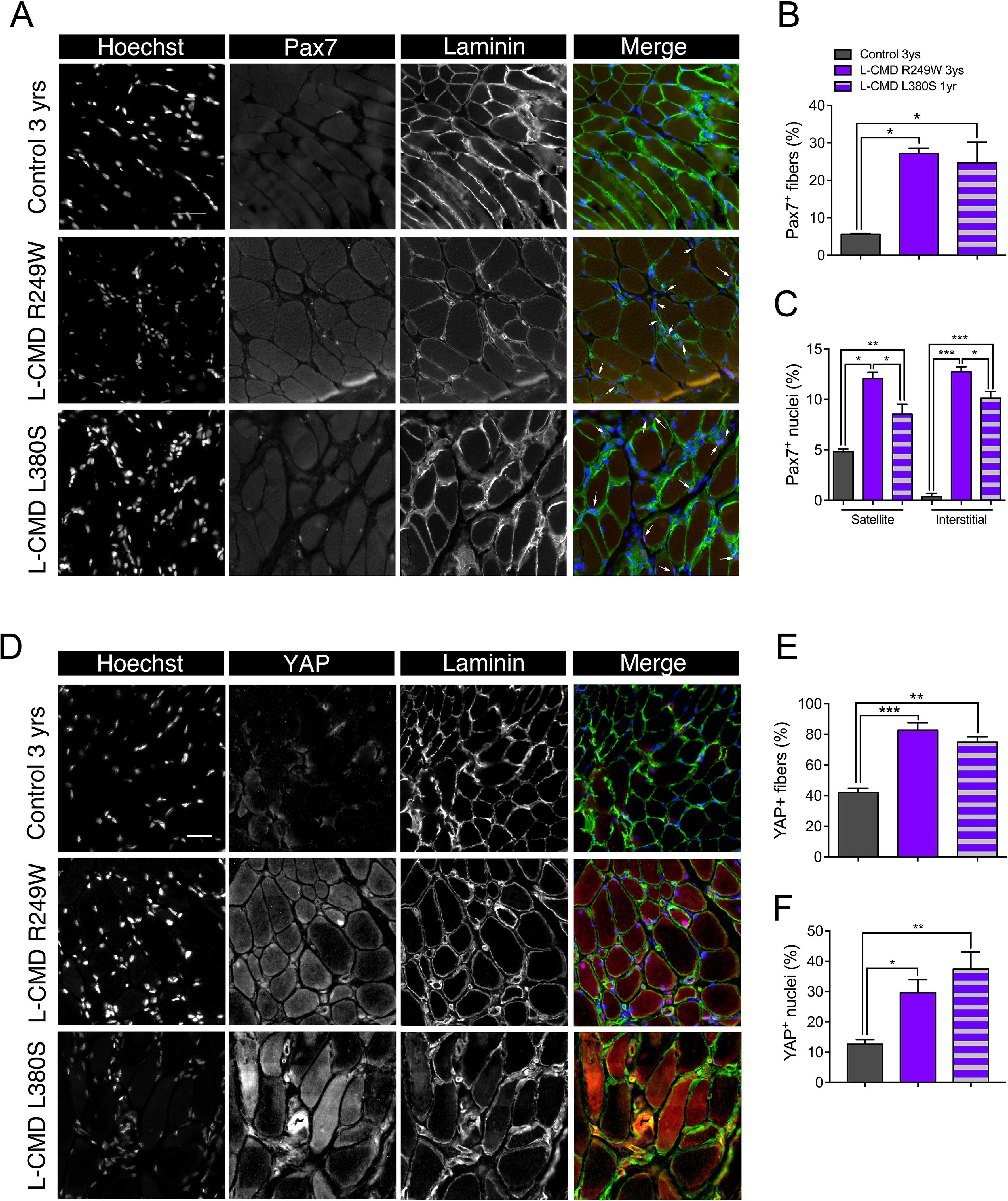
Histological data from muscle biopsies of patients with *LMNA*-CMD. (**A**) Immunofluorescence images of Pax7^+^ (green) and laminin (red) in muscle section from a control 3-year-old boy, a 3-year-old boy with R249W mutation and a 1-year-old boy with L380S mutation. Nuclei are stained with Hoechst (blue). Scale bar: 50 µm. (**B**, **C**) Quantification of Pax7^+^ cells per fiber and Pax7^+^ cells per nucleus in control and *LMNA*-CMD patients. Pax7^+^ cells in satellite or interstitial positions were determined. * p<0.05 versus control. (**D**) Immunofluorescence images of YAP (green) and laminin (red) in muscle section from a control 3-year-old boy, a 3-year-old boy with R249W mutation and a 1-year-old boy with L380S mutation. Nuclei are stained with Hoechst (blue). Scale bar: 30 µm. (**E, F**) Quantification YAP^+^ cells per fiber and YAP^+^ cells per nucleus in control and *LMNA*-CMD patients. * p<0.05 versus control. Values are expressed as means ± SEM

## DISCUSSION

A-type lamins are major nuclear proteins involved in mechanosensing and signaling between the nucleus, the cytoskeleton and the extracellular matrix. There is mounting evidence that lamins and nucleo-cytoskeletal coupling are required for cellular and nuclear mechanotransduction, muscle development and plasticity. The mechanisms by which *LMNA* mutations result in muscle-specific defects remain unclear, thus preventing the development of effective therapeutic approaches. In a recent study, nuclear defects including nuclear envelope rupture, DNA damage and chromatin protrusions have been correlated with the severity of muscle laminopathies [13]. It remains unclear how nuclear defects impair muscle differentiation and growth. Here, we describe a process by which *LMNA* mutations responsible for congenital muscle dystrophy impaired the ability of skeletal muscle to hypertrophy in response to a mechanical challenge, thus altering muscle growth. This relies on disorganized adhesion junctions between MuSCs, aberrant YAP signaling and fusion defects. Thus, our data highlight a critical role of A-type lamins in modulating skeletal muscle growth.

Mechanical stimuli are transferred to the actin cytoskeleton, leading to the activation of various signaling pathways that alter cellular dynamics and ultimately control key cell fate decisions such as proliferation and growth arrest. The transcriptional co-activator YAP is tightly regulated by the actin cytoskeleton and has been implicated as a main signaling protein in skeletal muscle mechanotransduction [38]. In activated satellite cells and proliferating MuSCs, YAP is predominantly nuclear permitting cell proliferation [33, 39]. In a differentiated post-mitotic multicellular context, it is the physical and architectural properties of the cellular microenvironment that inform the cell of its proliferative capability, a process that is controlled by YAP/TAZ signaling [40]. Indeed, knockdown of YAP results in impaired proliferation presumably by desensitizing the cell to its physical constraints [41]. As MuSCs exit the cell cycle and fuse to form multinucleated myotubes, YAP is phosphorylated by LATS 1/2 kinase and sequestered in the cytoplasm by 14-3-3 proteins, rendering YAP inactive [42]. However, YAP can be reactivated in myotubes by mechanically stretching cells [40] and in adult myofibers by functional overload of muscle [36].

There is growing evidence that A-type lamins are required for normal YAP signaling. We have previously shown that human derived A-type lamin mutant myoblasts are unable to traffic YAP to the nucleus in response to cyclic strain [14]. Others have demonstrated by traction force mapping that force transfer from the cytoskeleton to the nucleus is dependent on the LINC complex proteins and is critical for YAP trafficking and transcriptional activity [43]. We show here that mechanically stretching human primary myotubes results in YAP translocation to the nucleus, whereas laminopathic (ΔK32) mutant myotubes showed an aberrant, mirrored response. In *LMNA* ΔK32 myotubes, YAP was nucleoplasmic in static conditions and was exported from the nucleus following cyclic stretch with a concomitant decrease in the N/C ratio (Fig. 4). This YAP signaling defect was accompanied by a lack of actin cytoskeleton remodeling that was observed in WT myotubes following stretch (Fig. 4). Moreover, WT mice showed a similar response *in vivo*, with increased YAP^+^ fibers following functional overload whereas *Lmna*^+/ΔK32^ control mice had more YAP^+^ fibers, which decreased following functional overload. Importantly, we show that these defects appear to be specific to congenital laminopathies, as the LmnaH222P/H222P mouse model of EDMD was able to respond to FO comparably to WT mice (Supp. Fig. 1).

A key finding from our study was that muscle progenitor fusion defects are present in *LMNA*-CMD muscle derived cells and a mouse model carrying an A-type lamin mutation responsible for *LMNA*-CMD. Human muscle cross sections from patients with *LMNA*-CMD have a greater number of Pax7^+^ cells, thus supporting increased proliferation of MuSCs and/or defective incorporation of activated MuSCs into the myofibers. By first implementing *in vitro* experimentation, we demonstrate that fusion defects exist in *LMNA*-CMD cells and may be due to aberrant adherens junction formation. Adherens junctions are large macromolecular complexes that accumulate at cell-cell contacts, the formation of which requires cadherins and catenins [44–47]. In our study, M-cadherin and β-catenin were poorly organized in confluent *LMNA*-CMD MuSCs, whereas confluent WT counterparts displayed a typical zipper-like formation, characteristic of force-bearing adherens junctions [48]. Importantly, adherens junctions are crucial for cellular mechanonosensitivity, permitting mechanical forces to mediate cellular behavior. In skeletal muscle, cadherin-mediated adhesions contribute to the quiescence of MuSCs in the niche cells by providing structural integrity, mechanosensation, cell polarity and juxtacrine signalling [49, 50]. Disruption of cadherin-based adhesion between MuSCs and the myofiber is a critical step allowing the transition from quiescence to activation and proliferation [50]. One likely mechanism of mechanical activation of MuSCs is that mechanical stretch on cadherin-based adhesions alter physically and/or functionally cadherins, thus leading to the departure of MuSCs from the niche, and activation [49, 50]. Altered cadherin-based adhesions and/or impaired mechanosensing may in turn promote activation and proliferation of MuSCs, contributing to the higher number of Pax7^+^ cells present in *LMNA*-CMD patients (Fig. 8). Reduced M-cadherin expression in the current study may be another key factor contributing to the impaired differentiation and plasticity of A-type lamin mutant muscle. Numerous *in vitro* studies have shown that M-cadherin is essential for the differentiation of myoblasts into myotubes [44, 51–53]. Although M-cadherin is necessary for myoblast fusion it does not appear to affect the induction of myogenesis [54], which supports our findings that A-type lamin mutant myoblasts were able to induce myogenin protein expression but displayed a lower fusion index than wild type MuSCs (Fig. 1). Overall, the data presented here support our proposition that M-cadherin-β-catenin complexes may be affected by dysregulation of actin remodelling proteins that subsequently leads to defective YAP localization and impairments in muscle differentiation and plasticity.

Adaptation of skeletal muscle to physical challenges is accompanied by NMJ remodelling, such that both endurance and resistance exercise cause phenotypic changes in pre- and post-synaptic structures [55]. Accordingly, in our study WT and to a limited extent *Lmna*^+/ΔK32^ NMJs increased in size without any change in pre-post synaptic overlap following FO (Fig. 7). However, whereas WT NMJs in overloaded muscle displayed a typical continuous pretzel-like structure, the postsynaptic network of overloaded *Lmna*^+/ΔK32^ mice was highly fragmented suggesting a compromised maintenance of NMJs. The molecular mechanisms by which NMJ plasticity is compromised in lamin-mutated muscle may be attributed to YAP deregulation. YAP is a crucial regulator of neuromuscular junction formation and regeneration. In muscle-specific YAP mutant mice, postsynaptic and presynaptic differentiation and function was impaired and subsequently inhibited NMJ regeneration after nerve injury [56].

## Conclusions

We show here that functional A-type lamins are critical to allow fine tuning of the appropriate mechano-signaling required for skeletal muscle growth. *LMNA* mutations responsible for congenital muscle dystrophy impaired the ability of skeletal muscle to hypertrophy in response to a mechanical challenge due to impaired fusion of satellite cells, aberrant YAP signaling and impaired neuromuscular junction. Thus, our data highlight a critical role of A-type lamins in modulating skeletal muscle growth.

**Table 1.**
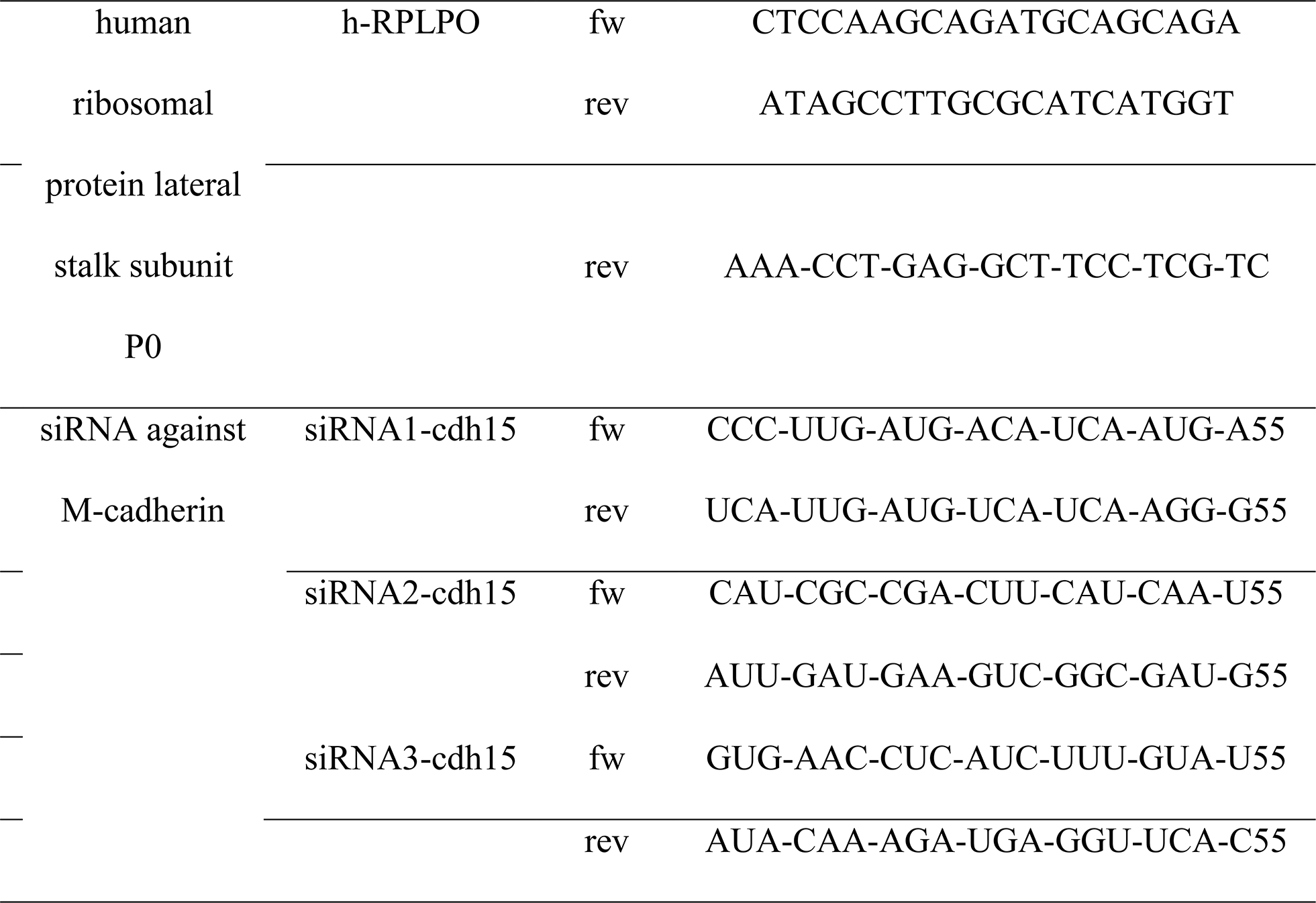
Primer and siRNA sequences

**Table 2.**
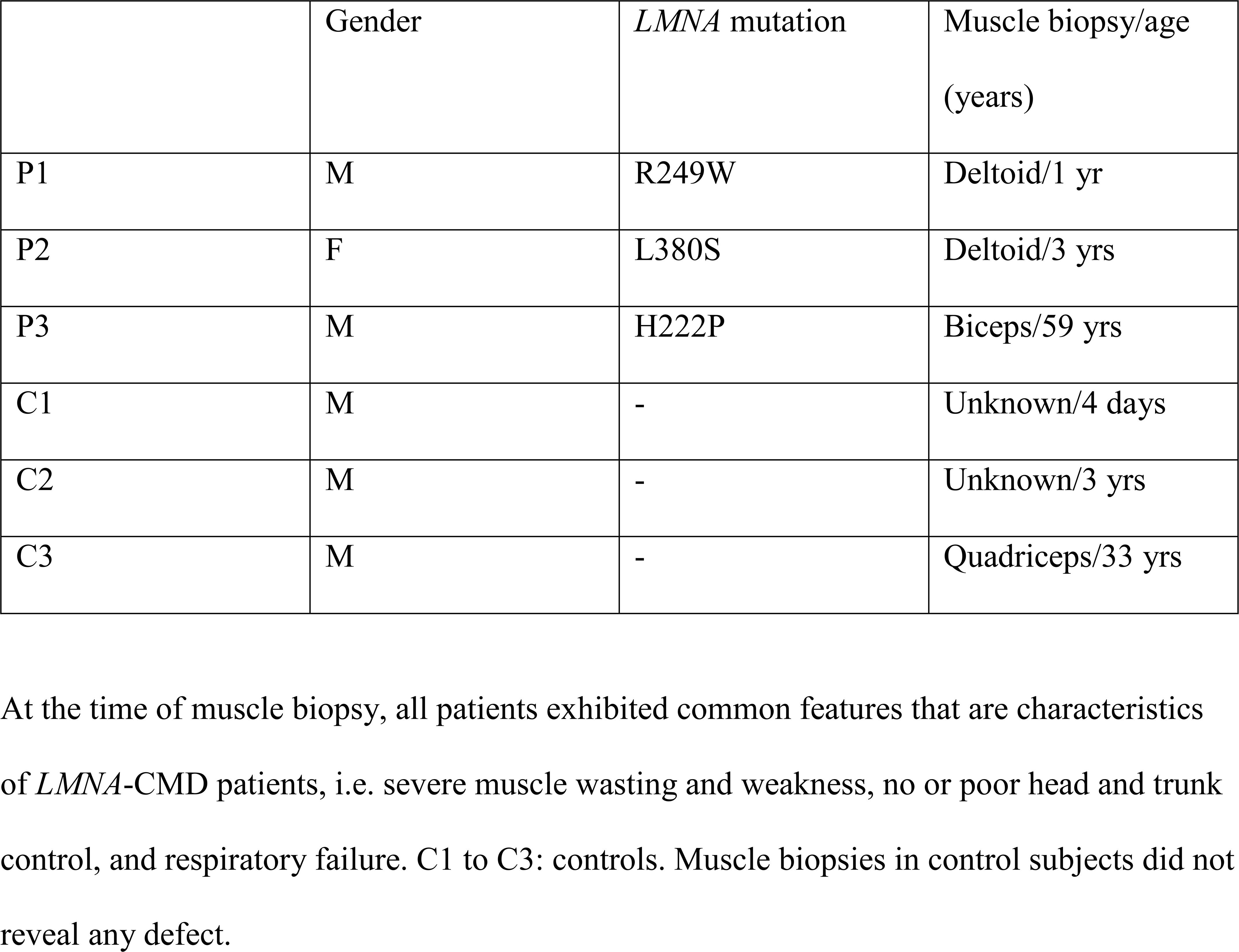
Characteristic of patients.

## Author contributions

Conceptualization, CC.; methodology, CC, DO..; validation, C.C. and DO..; formal analysis, C.C., DO., and NR.; investigation, C.C., DO., MV., and SM.; resources, K.M., AB., GB.; EL. and NR.; writing—original draft preparation, C.C. and DO; writing—review and editing, CC., DO., MV., GB., GBB., AB., NR., and EL.; supervision, CC.; project administration, CC. and GB.; funding acquisition, CC. and GB.

## Acknowledgments

We thank the IRIS-platform (Sorbonne University) for imaging facility. We also thank Patricia Davidson for constructive discussion. The authors certify that they comply with the ethical guidelines for publishing in the Journal of Cachexia, Sarcopenia and Muscle [57]

## List of abbreviations

AChR: Acetylcholine receptor
BSA: bovine serum albumin
EDMD: Emery-Dreifuss muscular dystrophy
ΔK32: *LMNA* c.94_96delAAG, p.Lys32del
hEGF: human epidermal growth factor
CMD: congenital muscular dystrophy
FO: functional overload
L380S: *LMNA* p.Leu380Ser
MuSC: muscle stem cell
NF: neurofilament
PBS: phosphate buffer solution
PLN: plantaris muscle
R249W: *LMNA* p.Arg249Trp
SDS: sodium dodecylsulfate
SRF: serum responsive factor
Syn: synaptophysin
TBS-T: tris-buffered saline-tween
WT: wild-type
YAP: Yes-Associated Protein

---

**Suppl Fig 1. Cell culture and plantaris muscle characteristics in *Lmna*^H222P^ mutation. (A)** Confocal immunofluorescence images of myosin (MF20, green) in WT and *LMNA*^H222P^ cells after 3 days of differentiation. Nuclei are stained with Hoechst (blue). Scale bar=50 µm. **(B)** Fusion index in WT and *LMNA*-CMD mutant cells after 3 days of differentiation. Values are expressed as means ± SEM. (**C**) Plantaris muscle mass normalized by body mass from WT and ***Lmna***^H222P^ mice in control and after 4-weeks FO. * p<0.05 versus control conditions. Values are expressed as means ± SEM

**Suppl Fig 2. Nuclear deformations in *Lmna*^+/ΔK32^ mice following functional overload. (A)** Confocal immunofluorescence images of nuclei (Hoechst, white) in WT and *Lmna*^+/ΔK32^ in control and after 7-days FO. (**B**) Nucleus deformations in WT and *Lmna*^+/ΔK32^ in control and after 7-days FO. Values are means ± SEM, n≥230 nuclei/condition.

**Suppl Fig 3. Histological data from muscle biopsies of controls and a patient with EDMD. (A)** Immunofluorescence images of Pax7^+^ (green) and laminin (red) in muscle section from a control 4-day-old boy, a control 33-year-old man and an EDMD patient 59-year-old with heterozygous *LMNA*^H222P^ mutation. Nuclei are stained with Hoechst (blue). Scale bar: 30 µm. **(B**) Immunofluorescence images of YAP (green) and laminin (red) in muscle section from section from a control 4-day-old boy, a control 33-year-old man and an EDMD patient 59-year-old with heterozygous *LMNA*^H222P^ mutation. Nuclei are stained with Hoechst (blue). Scale bar: 50 µm. (**E, F**) Quantification YAP^+^ cells per fiber and YAP^+^ cells per nucleus in controls and EDMD patient.

