## Supplementary figures and images for "Lamin-related congenital muscular dystrophy alters mechanical signaling and skeletal muscle growth"

### Suppl Fig 1

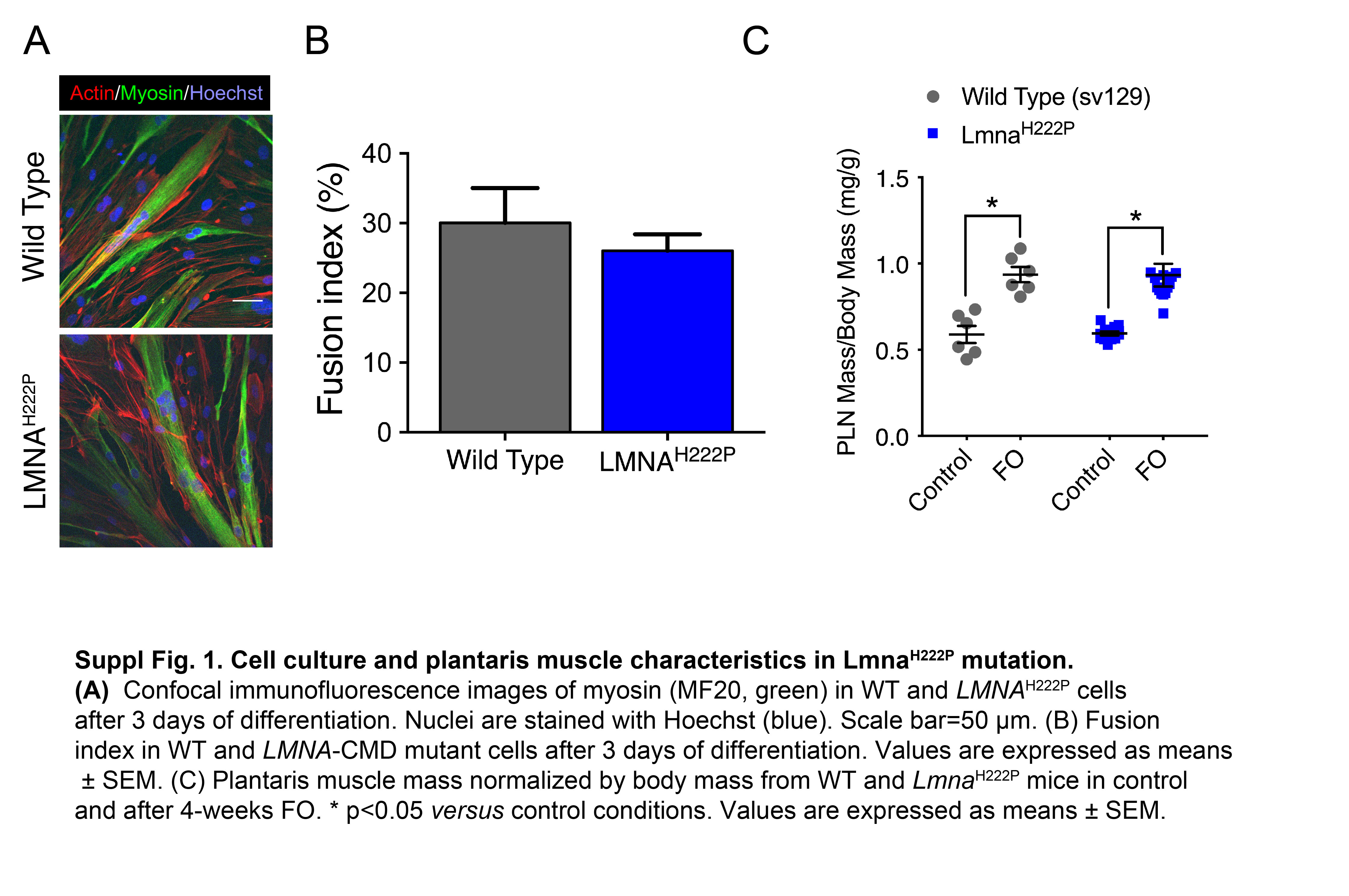

### Suppl Figure 2

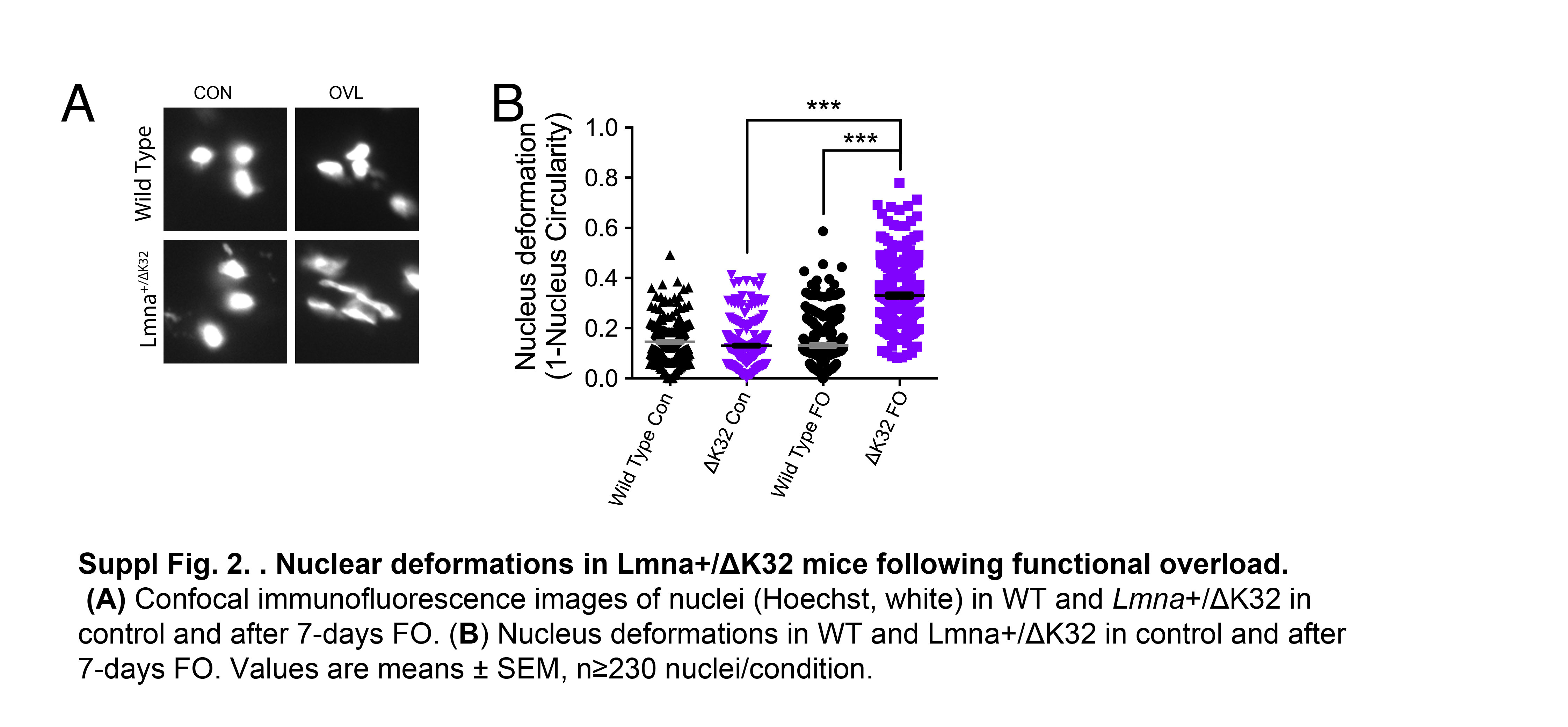

### Suppl Figure 3

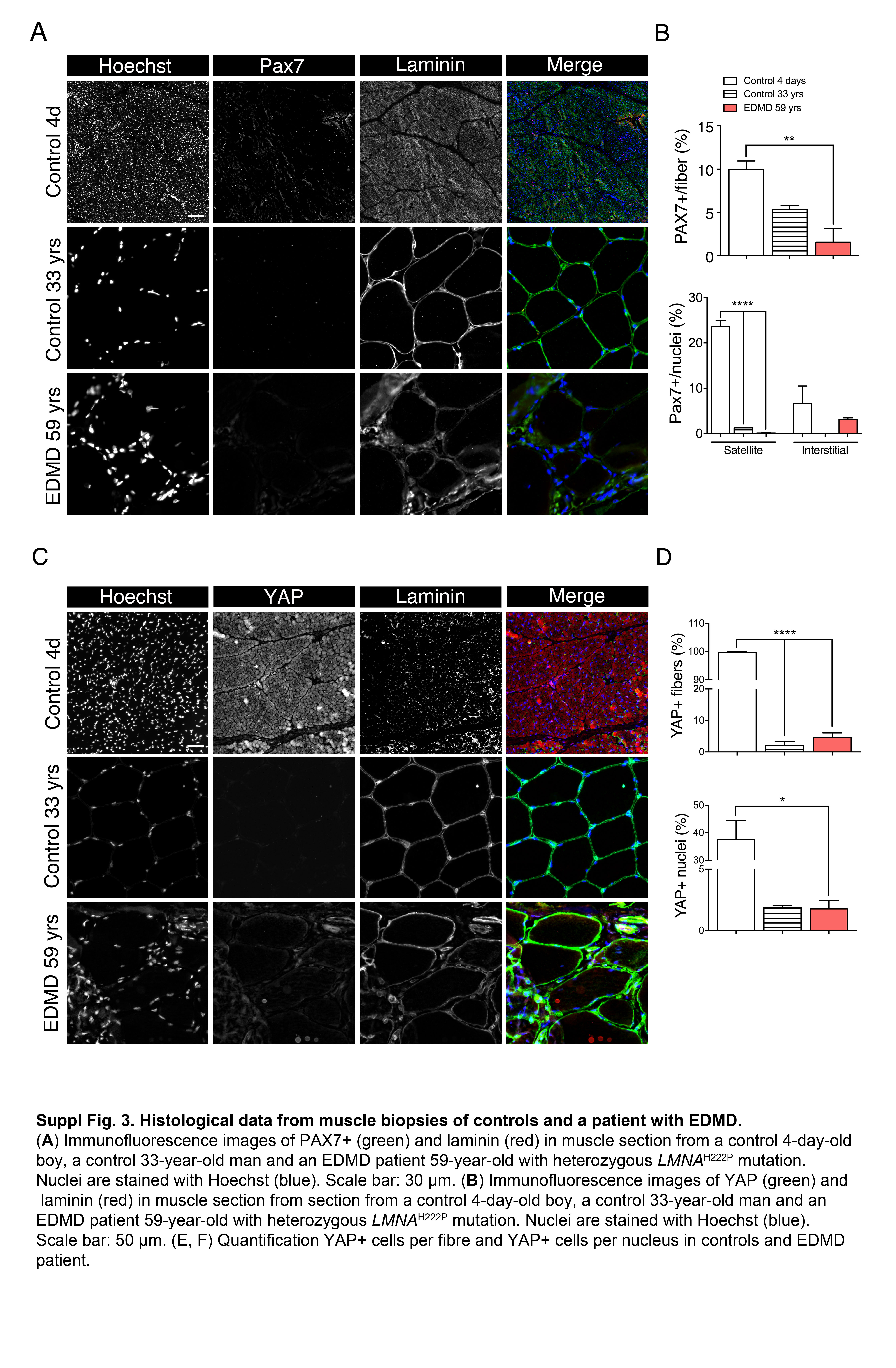
